# Unusual Morphological Changes of *Rugositalea oryzae*, A Novel Wrinkled Bacterium Isolated from The Rice Rhizosphere, Under Nutrient Stress

**DOI:** 10.1101/2022.08.29.505124

**Authors:** Young Ryun Chung, Jung Eun Lee, Zubair Aslam, Eu Jin Chung, Kwang Hee Lee, Byung Ho Kang, Ajmal Khan, Sarbjeet Niraula, Woo-Suk Chang

## Abstract

Bacterial cell morphology might result from natural selection to gain a competitive advantage under environmentally stressful conditions such as nutrient limitation. A bacterial strain YC6860^T^ isolated from the rhizosphere of rice (*Oryza sativa* L.) showed pleomorphic behavior with smooth cell morphology and wrinkled surface rods depending upon nutritional conditions. Based on scanning and transmission electron microscopy studies, we hypothesized that the surface-to-volume ratio of cells increases with decreasing nutrient concentrations. The transition from smooth to wrinkled cell surface morphology could be one of the adaptation strategies by which YC6860^T^ maximizes its ability to access available nutrients. To characterize the properties of the wrinkled strain, we performed taxonomic and phylogenetic analyses. 16S rRNA gene sequencing results showed that the strain represented a novel, deep-rooting lineage within the order *Rhizobiales* with the highest similarity of 94.2% to *Pseudorhodoplanes sinuspersici* RIPI 110^T^. Whole genome sequencing was also performed to characterize its genetic features. The strain YC6860^T^ might belong to a new genus, named *Rugositalea*, and a new species, named *Rugositalea oryzae*, In addition, taxonomic analysis showed that YC6860^T^ is Gram-negative, aerobic, and rod-shaped with large regular wrinkles resembling a delicate twist of fusilli, measuring 0.5-0.6 µm in width and 1.5-1.6 µm in length under nutrient-limiting conditions. This unique cell structure with regular rugosity could be the first finding that has not been reported in the existing bacterial morphology.

## Introduction

Bacteria have evolved over a long period of time to adapt to stressful environments by changing their shape and size. Different types of bacterial cell morphology, including rod, coccus, filament, star-like, helical/spiral and prosthecate shapes^1–5^, likely result from the evolution of bacteria and thus represent the basis of bacterial classification^6,7^. Morphological alterations of bacterial cells under environmental fluctuations in nutrient availability, temperature, space availability, and moisture content have been extensively studied^8–12^. Nutrient limitation occurs most often in an environment with active microbial competition. Under such adverse conditions, bacteria have evolved competitive mechanisms, by which they produce digestive enzymes or other protective and storable metabolites to outcompete their neighbors^13^. The slow-down of cell division, switching to fermentation, and alteration in cell morphology are also important metabolic and phenotypic strategies of survival^6,13^. The competition could induce morphological changes in bacteria such as curved, elongated stalk, branched and filamentous shapes to increase attachment to host surface or efficient nutrient uptake. The role of stalk lengthening in *Caulobacter crescentus* is to enhance nutrient absorption in oligotrophic aquatic environments with relatively low nutrient concentrations^8,14,15^. In addition, the altered cell size and shape to effectively improve nutrient uptake under nutrient deficiencies has been documented in model species such as *Escherichia coli* and *Bacillus subtilis*^2,7,16^.

Nutrient availability is the strongest factor affecting bacterial growth and survival in the rhizosphere. The rhizosphere is an energy-rich zone containing up to 40% of host photosynthetic carbon, and therefore microbial interactions in the rhizosphere are associated with various types of plant exudates^17^. The root exudates consisting of low-and high-molecular weight compounds may attract different microbial communities. Certain flavonoids are known to initiate the symbiotic process between legumes and nitrogen-fixing bacteria, while others attract beneficial microbes (e.g., plant growth-promoting rhizobacteria, PGPR) that could suppress harmful microbes or induce host resistance against plant pathogens or abiotic stresses^18,19^. In addition to flavonoids, microbial competition for nutrients within the rhizosphere could be triggered by the elevated level of microbial population, plant genotypes, and types of root exudates^20–22^.

In nutrient deficiency, the surface area-to-volume (S/V) ratio is critical because the nutrient uptake could depend on the exposed surface area. In order to achieve the large S/V ratio, bacterial cells may limit their size or transform their regular morphology to filamentous or prosthecate shapes^7^. For instance, the S/V ratio of rod-shaped bacterial cells remains constant under growth, although it decreases in spherical-shaped bacteria^23,24^. Peptidoglycans and cytoskeleton consisting of actin filaments and fatty acids are key factors in the morphological transition of bacterial cells^1,10,25^. In bacterial species, cell division begins with a coiled tubulin-like cell protein FtsZ which assembles the cell division machinery and interacts with the actin-like cytoplasmic proteins, resulting in a rod-shaped structure. The three types of actin proteins include MreB, Mb1 and MreBH. Among them, MreB plays a key role in the rod-shaped structure and movement of the daughter chromosome to the opposite pole. However, MreB is absent in the coccal species. The formation of MreB spirals depends on the presence of FtsZ in rod-shaped cell division^26,27^. In nutrient-rich media, the nutrients act upon metabolic sensors, which increase the cell size via FtsZ assembly inhibition, or increase the Z-period length before cytokinesis. In nutrient-poor media, the protein expression was decreased, randomly dispersed and had little effect on FtsZ assembly, resulting in reduction of the average cell size^1,16,28^. Recently, the mechanisms underlying the evolution of altered bacterial shape have been elucidated via the analysis of genome sequence data^2,29^. By mapping morphological phenotypes onto phylogenetic trees, we were able to gain an insight into how bacterial shapes evolved. Some morphologies such as helical and filamentous types, are spread all over the bacterial domain, indicating that they have evolutionally several origins^2^. However, the same morphology is clustered together in a specific region of the phylogenetic tree, suggesting that this morphology came from a shared ancestor^2^.

In this study, we hypothesized that the altered cell morphology from smooth to wrinkled rods of bacterial strain YC6860^T^ increased the S/V ratio of cells under nutritional stress. The hypothesis was proved by phenotypic observations of the strain cultivated under different nutrient concentrations using electron microscopy. Not only was the novelty of this strain characterized by a polyphasic approach, but also its full genome sequence was analyzed.

## Results

### Growth and morphology of the strain YC6860^T^

During the analysis of bacterial community in the rhizosphere of rice cultivated under conventional and no-tillage practices, the strain YC6860^T^ was isolated from the rice rhizosphere in the reproductive phase at a no-tillage paddy field^30^. Under SEM, the rod cells showed unique wrinkled features comprising grooves and margins uniformly distributed over the bacterial surface (Fig. 1a, b). However, the cells grown in 0.5 TSB for 7 days showed irregular rod or cocci shape without regular wrinkles, which we consider as smooth surface (Fig. 1c). Under TEM, the cell structures including wall and membrane were altered together in grooves and margins of the wrinkles (Fig. 1d, e, f). To confirm this unique change in cell shape, the strain YC6860^T^ was cultured in nutrient media at two different concentrations and the morphology was compared. The strain grew slowly in high-nutrient media (0.5 TSB) showing a normal growth curve until 15 days with increasing bacterial population and showed smooth shape of cells. While the cells grown in low-nutrient media (0.1 TSB) reached the steady state after 7 days of growth starting from 5 days post-inoculation and the cells were wrinkled (Fig. 1f, g). The morphology of this strain cultivated in 0.1 LB and 0.5 LB media was also similar in pattern to the strain under 0.1 TSB and 0.5 TSB (supplementary Fig. S1a, b). This result suggested that nutrient concentration of the culture media could be the main factor underlying the change in shape from smooth to the wrinkled form.

**Figure 1.**
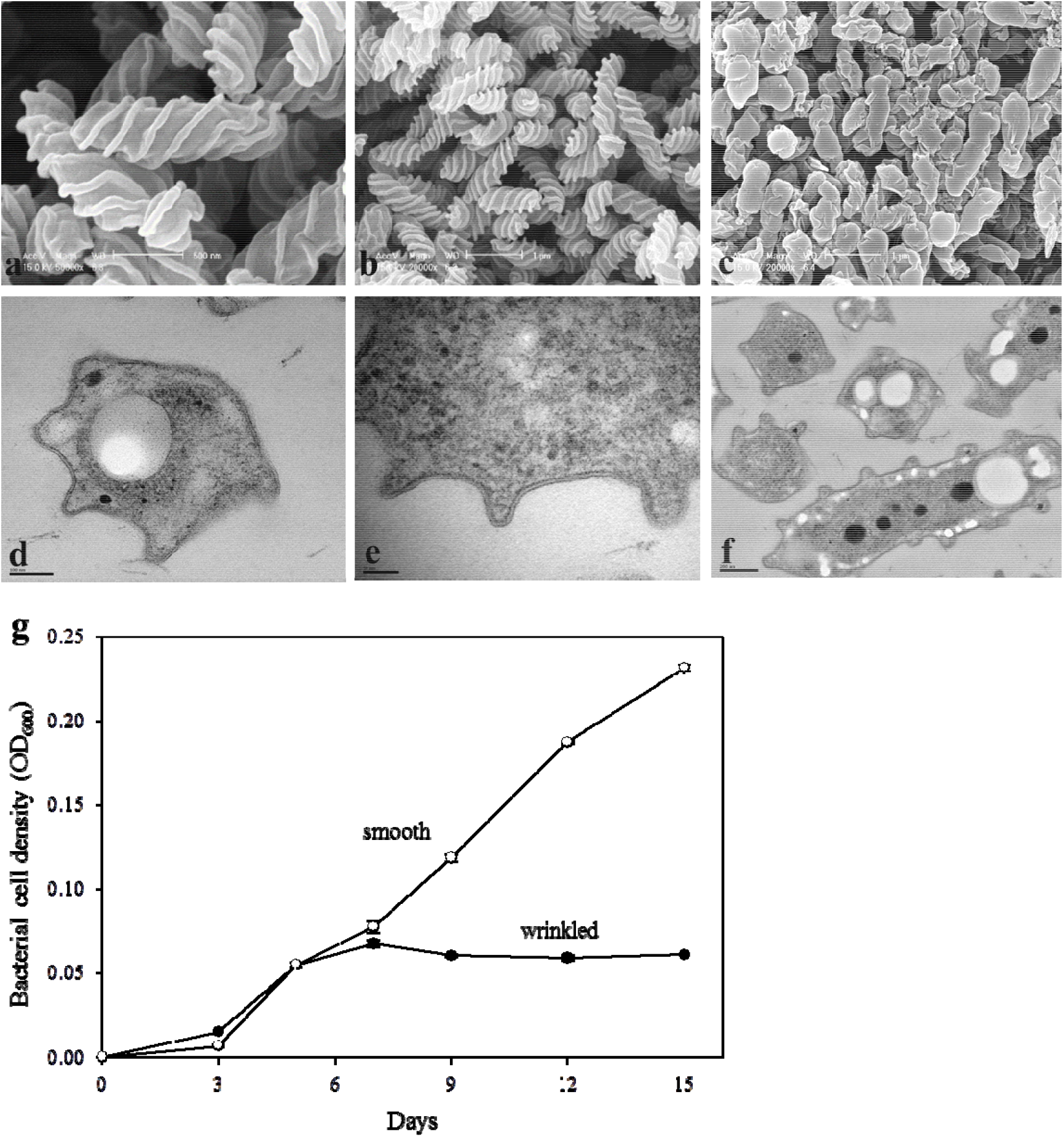
The effects of nutrient media on the growth of *Rugositalea oryzae* YC6860^T^. Bacterial cells grown in 0.1 TSB (a, b) and 0.5 TSB (c) on a rotary shaker at 28°C for 15 days were observed under SEM. Wrinkled cells were observed under TEM (d-f). Bacterial cell density in 0.1 TSB (closed circles) and 0.5 TSB (open circles) was observed at different time intervals. The open circles show bacterial growth in smooth cells, while the closed circles indicate bacterial growth in wrinkled cells with time (days).

### Transition from smooth to wrinkled shape

To confirm the change in cell shape from smooth to wrinkled form under nutrient stress, the strain YC6860^T^ was initially cultivated in 0.5 TSB for 15 days and the 15-day-old cells were transferred to 0.1 TSB after washing with 0.1 TSB twice by centrifugation. Before the transfer, the cells collected at 7 days after inoculation (S7) in 0.5 TSB showed a smooth surface (Fig. 2a, d). One day after transfer of 15-day-old cells from 0.5 TSB to 0.1 TSB, the smooth cells started to convert to wrinkled form (SW1) (Fig. 2b, e). More interestingly, the grooves were more prominent at 7 days after the transfer (SW7) (Fig. 2c, f). In addition to electron microscopy, the population density of strain YC6860^T^ and TOC in culture media were measured at several time points before and after the transfer of the cultures at 15 days. The bacterial population increased slowly until 15 days in 0.5 TSB, but increased rapidly for 3 days after transfer to 0.1 TSB. However, it started decreasing at 3 days after the transfer (Fig. 2g). TOC in culture media was also monitored at different time intervals to ensure transitions of nutritional levels. Indeed, the TOC in 0.5 TSB was not significantly changed until 15 days before transfer to 0.1 TSB at a range of 4,200~4,700 µg/mL. As expected, it was decreased to levels less than 800 µg/mL after transfer to 0.1 TSB and kept the same level until 7 days after the transfer (Fig. 2g). Not only does this result provide evidence of a decrease in the nutrient concentration, but it also indicates that the nutritional transition from high to low altered bacterial shape from smooth to wrinkled form. One possibility to explain this unique change in bacterial shape could be linked to increased S/V ratios of bacterial cells to absorb nutrients more efficiently for the survival of YC6860^T^ (Fig. 2g).

**Figure 2.**
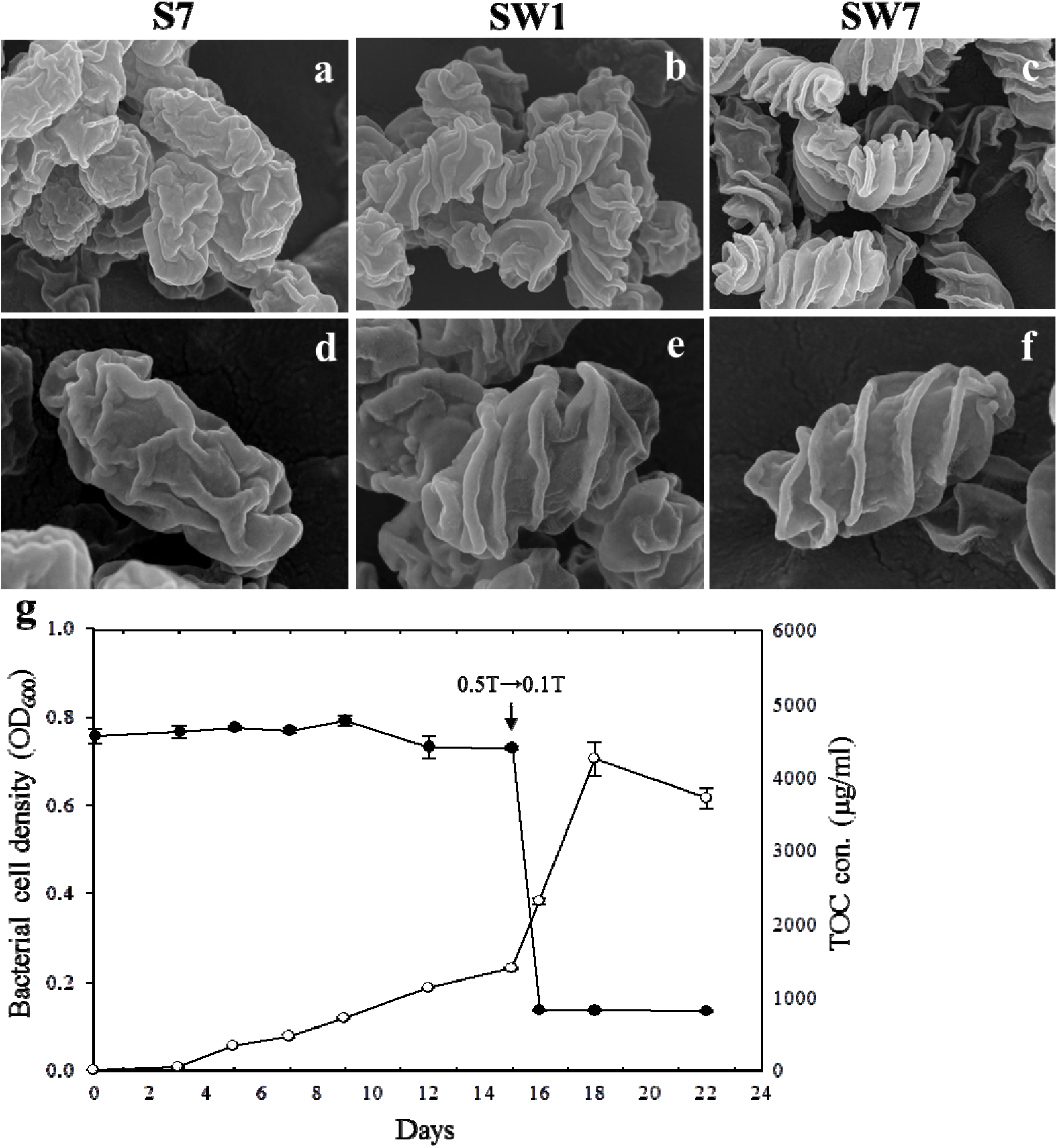
The effects of nutrient media on the growth of *Rugositalea oryzae* YC6860^T^. Bacterial cell shape was observed at 7 days after inoculation in 0.5 TSB (S7: a, d). The same cells were transferred to 0.1 TSB on day 15 after inoculation in 0.5 TSB and observed after 1 day (SW1: b, e) and 7 days (SW7: c, f) after transfer. The population density and the total organic carbon (TOC) were determined in cells growing in TSB media up to 22 days (g). The open circles show cell growth while the closed circles indicate TOC with time (days).

### Morphological transition correlated with surface area-to-volume ratio

The S/V ratio of a bacterial cell is related to its efficiency of nutrient uptake and respiration^7,24^. To confirm a possibility of changes in S/V ratios by transition from smooth to wrinkled form, the S/V ratio of YC6860^T^ cells was analyzed at five different nutrient concentrations ranging from 0.1 to 0.5 TSB. The randomly selected cells showed altered shape from wrinkled rods to less wrinkled and smaller cocci cells with the increase in nutrients (Fig. 3a). As shown in Figure 3b, c, the more nutrients, the less S/V ratios. The changes were statistically significant (*P* < 0.01) under TEM. As bacterial cells transit from smooth to wrinkled shape, they increase the S/V ratio, presumably to facilitate nutrient uptake. Regression analysis revealed strong inverse correlation between the nutrient concentration (X) and S/V ratio (Y) with the coefficient of determination r^2^=0.90, suggesting that with the decrease in each nutrient unit (e.g., 0.1X) in the range between 0.1 and 0.5 TSB, the S/V ratio of bacterial cells increased correspondingly during the transition (Fig. 3c).

**Figure 3.**
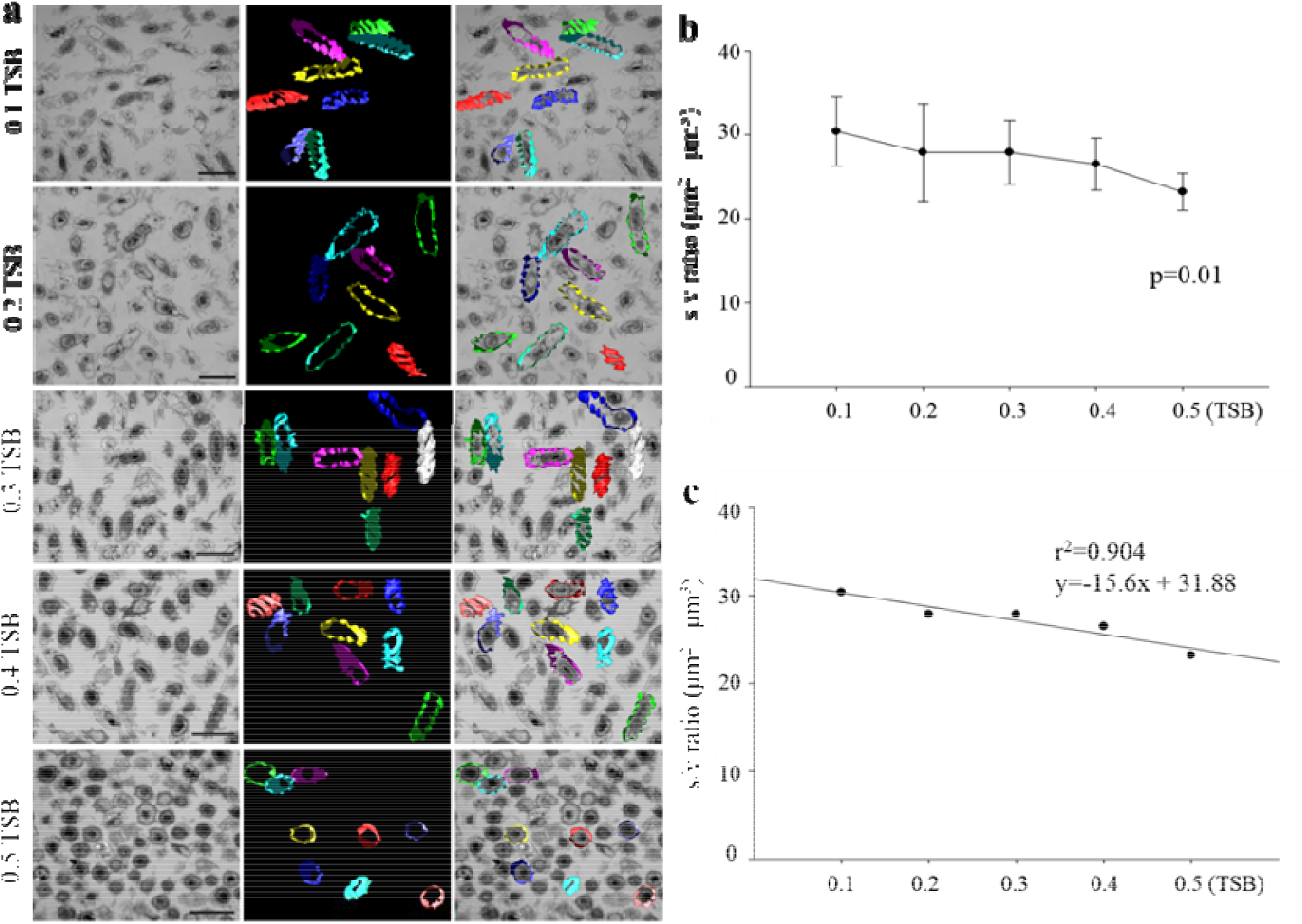
The surface-to-volume (S/V) ratio was determined in cells of *Rugositalea oryzae* YC6860^T^ grown in 0.1 to 0.5 TSB media for 15 days at 28°C using TEM (a). The decreased S/V ratio with increasing TSB concentrations; (b) and regression analysis (c). The second column of (a) shows three-dimensional models of cell surfaces; the models overlapped with bacterial cell images from the five TEM samples in the third column. Scale bars: 1 µm

### Phylogenetic analysis

The analysis of 16S rRNA gene sequence of strain YC6860^T^ (1,447 bp) compared with other sequences obtained from EzBioCloud server indicated that the strain belonged to order *Rhizobiales* of class *Alphaproteobacteria*. The phylogenetic tree constructed using a neighbor-joining method and maximum-parsimony using MEGA4 software showed that the strain YC6860^T^ was phylogenetically distinct from other related taxa (Fig. 4). The strain YC6860^T^ showed deep lineage with *Pseudorhodoplanes sinuspersici* RIPI 110^T^ and increased distance from *Rhodoplanes elegans* AS130^T^ with high bootstrap value. In addition, BLAST analysis revealed that the strain YC6860^T^ was most closely related to *P. sinuspersici* RIPI 110^T31^, *Rhodoplanes tepidamans* TUT3520^T32^ and *R. elegans* AS130^T33^ with a similarity of 94.2%, 93.7% and 93.7%, respectively. There was less than 93.0% similarity to other genera. Since the similarity is less than 95% at the genus level, we hypothesized that the strain YC6860^T^ could belong to a new genus.

**Figure 4.**
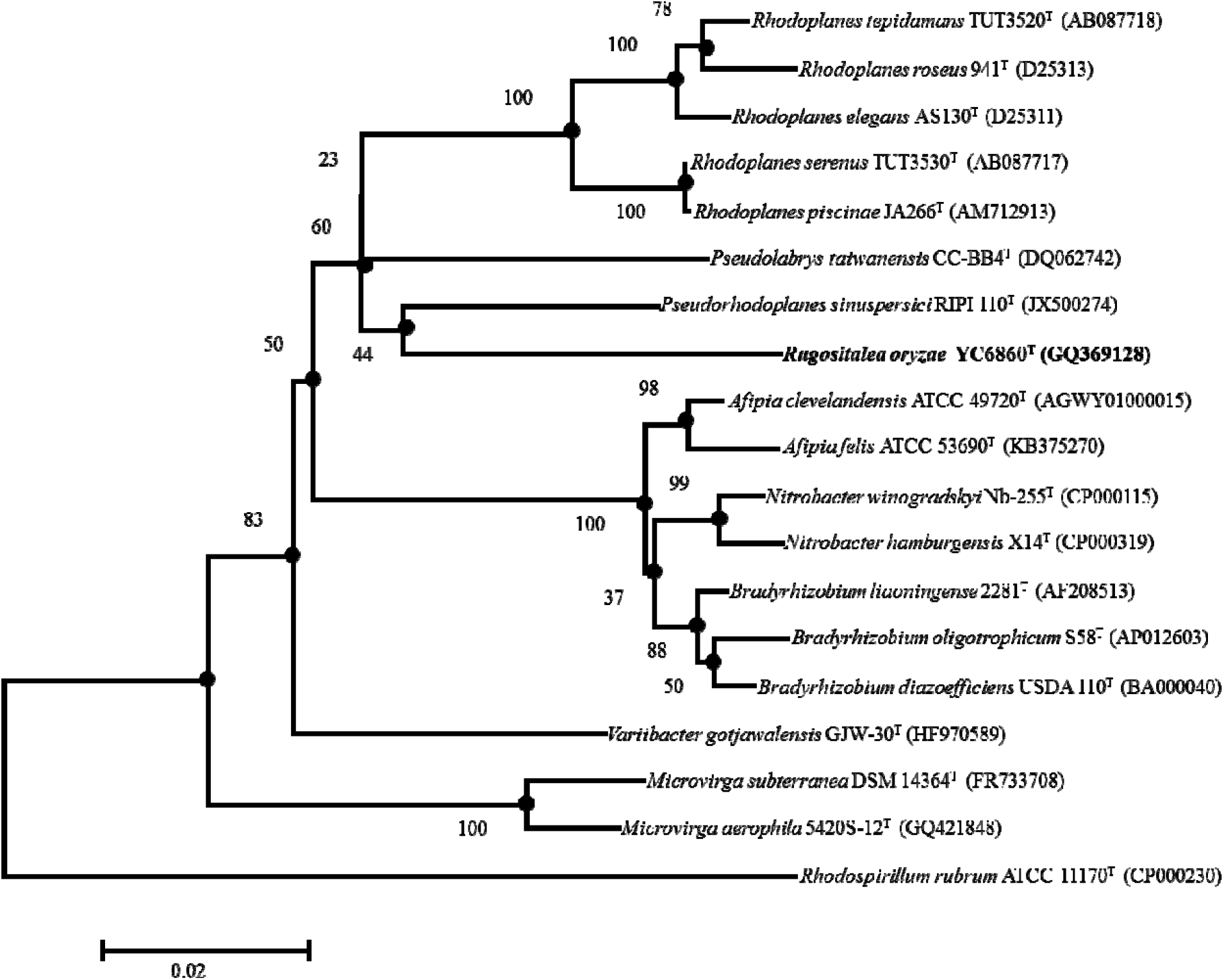
Phylogenetic tree reconstructed according to the comparative analysis of 16S rRNA gene sequences showing the relationships of *Rugositalea oryzae* YC6860^T^ and members of related genera. This phylogenetic tree was reconstructed using the neighbour-joining method and Jukes & Cantor evolutionary distance matrix data obtained from aligned nucleotides. Bootstrap values (expressed as percentages of 1000 replications) greater than 50% are shown at branch points. Filled circles indicate recovery of corresponding nodes in the maximum-likelihood and maximum-parsimony trees. Bar, 0.01 substitutions per nucleotide position.

### Phenotypic characterization

As strain YC6860^T^ is phylogenetically distinct from the closely related taxa based on 16S rRNA gene sequence similarity, other phenotypic traits such as morphological, biochemical and chemotaxonomic characteristics were compared with closely related genera. Unlike the strain YC6860^T^, *P. sinuspersici* RIPI 110^T^, *R. tepidamans* TUT3520^T^, *P. taiwanensis* CC-BB4^T^ and *R. elegans* AS130^T^ were isolated from oil contaminated soil, hot spring, soil and sludge, respectively^31–34^. Strains YC6860^T^ and *R. elegans* AS130^T^ are motile, but the others are non-motile. Salt is not necessary for their growth. The strains YC6860^T^ and *P. taiwanensis* CC-BB4^T^ are strictly aerobic, but the others are facultative anaerobes. The strain YC6860^T^ did not show any acid production or fermentation following reaction with the reagent in API 20E kit. In API ZYM kit tests, it showed esterase (C-4), leucine arylamidase, valine arylamidase, acid phosphatase, α-fucosidase, β-glucuronidase and α-mannosidase activities, and weak α-chymotrypsin, α-galactosidase, β-galactosidase, and N-acetyl-β-glucosaminidase activity. YC6860^T^ showed no alkaline phosphatase, lipase (C-14), or α-glucosidase activity, which differentiate it from the other strains. The DNA G+C content (63.5%) is different while major quinone (Q-10) remained the same as in other related strains (Table 1).

**Table 1.** Phenotypic variation in *Rugositalea oryzae* YC6860^T^ and phylogenetically related genera in the order *Rhizobiales*. Taxa: *Rugositalea oryzae* YC6860^T^; *Pseudorhodoplanes sinuspersici* RIPI 110^T^ (31); *Rhodoplanes tepidamans* TUT3520^T^ (32); *Rhodoplanes elegans* AS130^T^ (33); *Pseudolabrys taiwanensis* CC-BB4^T^ (34). ND, not determined; +, positive; −, negative.

| Characteristics | *Rugositalea oryzae* YC6860<sup>T</sup> | *Pseudorhodoplanes sinuspersici* RIPI 110<sup>T</sup> | *Rhodoplanes tepidamans* TUT3520<sup>T</sup> | *Rhodoplanes elegans* AS130<sup>T</sup> | *Pseudolabrys taiwanensis* CC-BB4<sup>T</sup> |
| --- | --- | --- | --- | --- | --- |
| Isolation source | Rice rhizosphere | Oil-contaminated Soil site | Hot spring | Sludge | Soil |
| Gram-staining | Negative | Negative | Negative | Negative | Negative |
| Motility | Motile | Non-motile | Motile | Motile | Non-motile |
| NaCl requirement | 0-1% | 0-3.5% | 0% | <1% | ND |
| Oxidase/ Catalase | +/+ | +/+ | ND | ND | +/+ |
| Temperature (°C) | 16-30 | 15-35 | 40 | 30-35 (optimal) | 15-36 |
| pH | 6.0-10.0 | 5.5-8.0 | 6.8-7.5 | 6-8.5 | 4.2-8.5 |
| Aerobic/ Anaerobic condition | +/- | +/+ | +/+ | +/+ | +/- |
| Alkaline phosphatase | - | ND | ND | + | + |
| Lipase (C14) | - | ND | ND | - | + |
| DNA G+C content (%) | 63.5 | 59.4 | 69.9 | 69.7 | 67.0 |
| Major quinone | Q-10 | Q-10 | Q-10, R-10 | Q-10, RQ-10 | ND |

\*Data for the related type strains are from this study except *P. sinuspersici* RIPI 110<sup>T</sup> (31) and *R. tepidamans* TUT3520<sup>T</sup> (32).

### Fatty acid analysis

The fatty acid profiles of YC6860^T^ and other closely related strains were compared. The fatty acids both C_16:0_ and C_18:1_ w7c were mainly detected in strains YC6860^T^, *P. sinuspersici* RIPI 110^T^, *R. tepidamans* TUT3520^T^ and *R. elegans* AS130^T^, but not in *P. taiwanensis* CC-BB4^T^ (Table S1). Interestingly, the main fatty acid C_15:0_ anteiso in CC-BB4^T^ was absent in the other strains including YC6860^T^. The fatty acid profiles of strain YC6860^T^ grown in 0.1 TSB and 0.5 TSB were also compared to examine the effects of different nutrient concentrations on the composition of major fatty acids. The cyclopropane fatty acid C_19:0_ cyclo w8c was ~3 times higher in cells cultivated in 0.1 TSB when compared with 0.5 TSB (Table 2). The saturated fatty acid C_10:0_ was detected only in cells in 0.5 TSB. However, the percentage of the main saturated and unsaturated fatty acids, C_16:0_ and C_18:1_ w7c, was not significantly different between the two conditions.

**Table 2.**
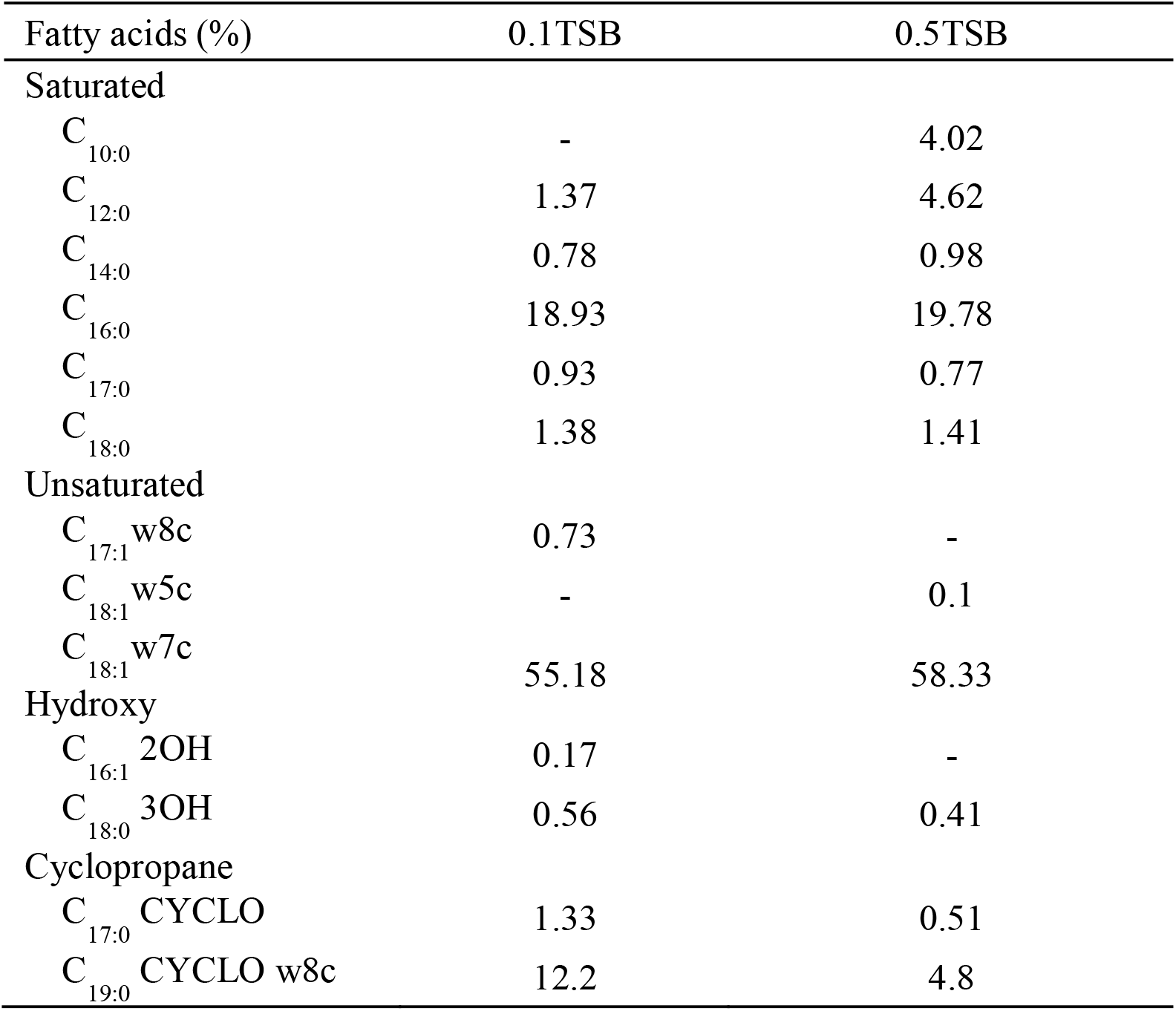
The cellular fatty acid profile of *Rugositalea oryzae* YC6860^T^ grown in TSB media at different nutrient concentrations. Data were expressed as percentages of fatty acid determined after cultivating the strain YC6860^T^ at 28°C for 10 days.

| Fatty acids (%) | 0.1TSB | 0.5TSB |
| --- | --- | --- |
| Saturated |  |  |
| C<sub>10:0</sub> | - | 4.02 |
| C<sub>12:0</sub> | 1.37 | 4.62 |
| C<sub>14:0</sub> | 0.78 | 0.98 |
| C<sub>16:0</sub> | 18.93 | 19.78 |
| C<sub>17:0</sub> | 0.93 | 0.77 |
| C<sub>18:0</sub> | 1.38 | 1.41 |
| Unsaturated |  |  |
| C<sub>17:1</sub> w8c | 0.73 | - |
| C<sub>18:1</sub> w5c | - | 0.1 |
| C<sub>18:1</sub> w7c | 55.18 | 58.33 |
| Hydroxy |  |  |
| C<sub>16:1</sub> 2OH | 0.17 | - |
| C<sub>18:0</sub> 3OH | 0.56 | 0.41 |
| Cyclopropane |  |  |
| C<sub>17:0</sub> CYCLO | 1.33 | 0.51 |
| C<sub>19:0</sub> CYCLO w8c | 12.2 | 4.8 |

### Genome description

The low 16S rRNA gene sequence similarity, phylogenetic analysis, and other phenotypic characteristics clearly distinguish the strain YC6860^T^ from all the species in the closely related genera *Pseudorhodoplanes, Rhodoplanes* and *Pseudolabrys*. Thus, we conclude that the strain YC6860^T^ is a novel genus with a novel species in the order of *Rhizobiales* with a proposed name *Rugositalea oryzae* YC6860^T^. The genome of *R. oryzae* YC6860^T^ comprises a single chromosome and is 8,193,889 bp long with a G□+□C content of 63.5% (5,205,286 bp) (Fig. S2). It is composed of a single scaffold and 12 island regions (Fig. S3). In total, 7,203,497 bp (87.91%) represent the coding capacity, and 7,776 genes were identified in the genome including 7,708 (99.13%) protein-coding genes. Among the protein-coding genes identified, the functional annotation of 6,069 (78.05%) genes was accomplished using different functional databases. Furthermore, 5,461 (70.23%) genes matched at least a single sequence in Clusters of Orthologous Groups database (COGs). The genes are distributed into COGs functional categories (Fig. S4). Protein-coding genes connected with KEGG Orthology (KO) were 3,317 (42.66%), and those connected with KEGG pathways and MetaCyc pathways were 2048 (26.34%) and 1585 (20.38%), respectively. The genome encodes 68 predicted RNAs including 3 rRNAs, 48 tRNAs and 17 other RNAs. No CRISPRs were found in the genome of YC6860^T^ using the CRISPRfinder. While 1290 (16.59%) coding genes were identified with signal peptides, 1,745 (22.44%) genes coded transmembrane proteins. Nucleotide and gene counts of the *R. oryzae* YC6860^T^ genome are summarized in supplementary Table S2.

## Discussion

Morphological changes in the cell often observed in diverse bacterial groups facilitate efficient nutrient uptake and enhance survival under stressful conditions such as low-nutrient levels. The shape and size vary with environmental conditions of bacterial transition from high to low nutrient levels^7,15^. The novel strain YC6860^T^ was isolated from the rhizosphere of rice, with fluctuating amounts of root exudates, which affect bacterial community and growth under different tillage systems^30^. The strain was rod-shaped with small irregular wrinkles in general media such as 0.5 TSA or 0.5 LB initially. However, when the strain was grown in more diluted low-nutrient media, its cell surface morphology changed from small irregular wrinkles to large regular wrinkles. Due to the complex natural conditions of the rhizosphere in terms of nutrient composition and microbial interactions, we used sterilized defined media that simulated different nutrient concentrations and negated microbial competition in the rhizosphere to investigate the effect of nutrients on the morphological changes in this strain^6,35^. The strain YC6860^T^ grew slowly in all the tested media compared with other commonly isolated bacteria from the rhizosphere. In 0.5 TSB, it showed a normal growth curve until 15 days with most of the smooth surface cells displaying irregular form. However, in 0.1 TSB its growth reached a stationary phase after 7 days with wrinkled cell surface, which might be attributed to nutrient limitation. Bacterial strains such as *Actinomyces israelis, Arthrobacter globiformis, Clostridium welchii* and a few *Pseudomonas* species were known to form the filamentous structure in nutrient-deficient media^7^. Similarly a bacterial species, *Caulobacter crescentus* in low phosphate media showed stalk elongation due to the presence of a high-affinity phosphate binding protein (Psts) for phosphate uptake and hydrolysis^36^. For confirmation of the shape change of this strain under nutrient limitation, the cells cultivated in 0.5 TSB were transferred to 0.1 TSB. The shape of the cells was gradually changed from fewer wrinkles to more conspicuous wrinkle formation with time, which was induced by decreased nutrient concentration. During the culture under different nutrient concentrations, the altered cell shape could reflect the adoptive behavior of the strain YC6860^T^ under nutritional stress as all other factors remained constant^2,7^. The bacterial cell density in the low-nutrient media decreased with time and also the TOC was reduced. The low nutrient levels slowed down the cell division and decreased the average cell size^16^. Conversely, the cell size of strain YC6860^T^ in 0.1 TSB was bigger than in 0.5 TSB, which might represent a stress factor for the growth of this strain. Instead, it developed prominent regular wrinkles on the cell surface uniformly in low-nutrient condition showing high S/V ratio. We therefore investigated the relationship between nutrient concentrations and S/V ratios using TEM. Results suggested a significantly inverse relationship between the rugoses of the cells and nutrient concentrations. One of strategies of the bacterial species to cope with nutrient limiting conditions is to increase S/V ratio for better access to available nutrients^7^. *C. crescentus* is the most extensively studied bacterium with elongated stalk, which increases the cell surface area but with little effect on the S/V ratio of the cell^14,29^. TEM analysis showed that rugoses occurred throughout the cell structure including inner and outer membrane of strain YC6860^T^ that increased the S/V ratio of the cells with decreasing nutrient concentration, which resulted in an increase in the space available for nutrient absorption under nutrient limiting condition.

On the basis of 16S rRNA gene sequence analysis, the strain YC6860^T^ was found to belong to the order *Rhizobiales*, which is a phenotypically heterogeneous assemblage of Gram-negative bacteria^37^. As the strain showed low sequence similarity (<94.3%) to the closely related members of genera *Pseudorhodoplanes, Rhodoplanes* and *Pseudolabrys*, it should be assigned a novel genus, with a name proposed as *Rugositalea* and a representative species *R. oryzae*. The physiological and biochemical characteristics of this strain also showed its distinct features from other related species. The fatty acid profile showed a high percentage of saturated fatty acid (C_16:0_) and unsaturated fatty acid (C_18:1_ w7c) levels in YC6860^T^, *Pseudorhodoplanes* and *R. elegans* AS130, but not in *P. taiwanensis* CC-BB4. The fatty acid composition of bacterial species is modified by the environmental changes especially pH, temperature and nutrient of growth media^38^. The alteration of cell size due to nutrient limitation is also caused by fatty acid biosynthesis, which controls cell expansion and permeability^25^. The fatty acid profile in strain YC6860^T^ at low nutrient media was distinct when compared with high nutrient media suggesting that the strain also responds to low-nutrient conditions by changing the fatty acid composition. However, the difference between fatty acid compositions at high and low nutrient media is not striking. The full genome of this novel strain was sequenced and deposited. As the strain YC6860^T^ exhibits extraordinary shapes depending on nutrient concentrations, the genome data might provide an opportunity to study the presence of unique metabolic pathways previously unknown for morphological evolution of bacteria^29^.

This study demonstrated only the influence of nutrients on the cell shape of the pleomorphic strain YC6860^T^ comprising smooth and wrinkled cells. There was decrease in the growth rate under low levels of nutrients due to reduced cell division but increase in cell size especially cell length in this strain. The rod length and development of large regular wrinkles in low nutrients showed the increased S/V ratio, indicating that bacteria manipulated the morphology to their advantage in a given environment^7,16^. Surprisingly, star-shaped cells similar to the wrinkled cell of strain YC6860^T^ were previously observed in the concentrated cell fluid of natural soil by TEM^39^. It was discussed that it could not be ruled out the possibility of an authentic form for certain soil microorganisms. It is not known how rugoses of the cell are formed in response to nutrient limitation. The actual mechanism underlying the evolution of shape in bacteria is not entirely clear; however, several shape-determining proteins such as actin-like cytoplasmic proteins, scaffold proteins and peptidoglycan are thought to determine the shape and size^27,40,41^. The mechanism underlying alterations in cell shape in this strain needs to be elucidated with further investigations into the molecular ecology.

*Rugositalea* (Ru.go’si.tal′e.a. L. adj. *rugosus* of rugose or wrinkled; L. fem. n. *talea* a rod; N.L. fem. n. *Rugositalea* a rugose forming rod) cells are aerobic, Gram-negative and motile with a polar flagellum (Fig. S5). The morphology is pleomorphic ranging from rod to cocci under different nutrient concentrations. Under low-nutrient conditions, the cells changed from rod to wrinkled shape with rugoses on the cell surface, reflecting restructured cell wall and membrane. The cells tested are positive for oxidase and catalase. The respiratory quinone is ubiquinone Q-10. The major fatty acids are C_18:1_ w7c and C_16:0_. *Rugositalea* belongs to the *Bradyrhizobiaceae* family within the order *Rhizobiales*. The type species is *Rugositalea oryzae*. The main features of *Rugositalea oryzae* YC6860^T^ (o.ry′za.e. L. fem. n. *Oryza* genus name of rice; L. gen. n. *oryzae* of rice, refers to the isolation of type strain from a rice field) are similar to those described for the genus: Creamy, smooth, flat and circular colonies, 0.1-0.5 mm in diameter and with entire margins, formed on 0.1 TSA after 30 days at 28°C. Cells measure 0.4-0.6 **×** 1.1-1.9 µm, dividing by binary fission in the mid-exponential phase. The temperature range for growth is 16-30°C (optimum 28°C). The growth pH range is 6-10 (optimum 7). It does not require NaCl for growth, but cannot tolerate NaCl above 0.5% (w/v). It exhibits esterase (C-4), leucine arylamidase, acid phosphatase, napthol-AS-BI-phosphohydrolase, α-fucosidase, valine arylamidase, β-glucuronidase, and α-mannosidase activities, and weak activity of β-galactosidase, β-glucosidase, α-chymotrypsin, α-galatosidase and *N*-acetyl-β-glucosaminidase. Alkaline phosphatase, lipase (C14) and α-glucosidase are negative. It grows on minimal agar medium supplemented with a small amount (0.3%, w/v) of inositol, melibiose, lactose, salicine, raffinose, glucose, sucrose and mannose as a single carbon source. However, it cannot grow in media containing fructose, mannitol and arabinose. It hydrolyzed hippurate but not chitin, casein or xylan. Nitrate is reduced to nitrite or nitrogen. The G+C content of the type strain is 63.5 mol%. The type strain, YC6860^T^ (=KCCM 43037^T^ =NBRC^T^ 109316), was isolated from the rhizosphere of no-tilled rice field at Jinju, Korea.

## Methods

### Isolation of strain

Soil samples for bacterial isolation were collected from the rhizosphere of rice paddy fields managed under no-tillage practices for 5 years, located at Gyeongsang National University farm (Daegok valley, 35° 14’ 21” N, 128° 13’ 23” E), Jinju, Korea. To isolate bacteria from the samples, one gram of soil was added to 10 mL buffer solution (50 mM phosphate buffer, pH 7.0) and half of the soil suspension was sonicated for 15 s with an electronic homogenizer (Bandelin Sonoplus, Berlin, Germany) at 260 W/cm^2^. Serial dilutions were obtained after mixing both sonicated and non-sonicated portions and 100 µl diluted (10^−3^-10^−5^) aliquots were spread on half-strength (0.5) R2A agar plates in large petri dishes (150 mm in diameter). These agar media supplemented with amphotericin B (50 µg/mL) to inhibit fungal growth, and 40% (w/v) soil extract were incubated at 28°C for more than a month. The small (less than 1 mm) bacterial colonies were selected on the basis of morphology and the isolates were purified and sub-cultured on 0.5 R2A. A strain YC6860^T^ was isolated and purified from the rhizosphere of no-tillage (Z2) rice field and stored at −70°C in 20% (w/v) glycerol stocks for further tests^30^.

### Bacterial culture preparation

Strain YC6860^T^ was cultured in various culture media with different nutrient composition such as tryptic soy broth (TSB), tryptic soy agar (TSA), R2A, LB broth (LB) and nutrient agar (NA) to determine the optimal medium for growth. All these media were purchased from Difco and diluted from one-tenth (0.1) to half-strength (0.5) for bacterial growth. During the first set of experiments, the growth of strain was better in 0.1 (3.0 g/L) ~ 0.5 (15 g/L) TSA or TSB media compared with other media, and therefore, selected for further experiments. Growth was measured by inoculating two to three loops of colonies developed on half-strength (0.5X) R2A agar plate for one month at 28°C into 100 mL 0.1 ~ 0.5 TSB media in 250 mL flasks, and incubated at 28°C with shaking (140 rpm) for the indicated periods. The optical density of a bacterial culture at 600 nm (OD_600_) was measured using a spectrophotometer (X-ma 1200, Human Corporation). The cells cultivated in 0.1 ~ 0.5 TSB media were collected by centrifugation and prepared for microscopic observation. Bacterial cells grown for 15 days on 0.1 TSA were characterized biochemically and physiologically.

### Bacterial cell morphology

Cell morphology of strain YC6860^T^ was observed under a Nikon light microscope at 1000x after Gram staining. The motility was determined using cells grown on 0.5 R2A agar plate for seven days at 28°C by phase-contrast microscopy. Flagella were observed via transmission electron microscopy (TEM) (Jeol, JEM 2010) after negative staining of specimens with 2% phosphotungstic acid on 200 mesh formvar-coated copper grids. The ultrathin sections of cells prepared using an ultramicrotome (Leica EM UC6) were double-stained with 2.0% uranyl acetate and lead citrate solution and used for ultrastructural observations. Scanning electron microscopy (SEM) was performed using bacterial cells cultivated in different culture broths at different concentrations (0.1 ~ 0.5 TSB) and recovered by centrifugation (8000*g* at 4°C) for 10 min and washed twice with 0.1 M phosphate buffered saline (PBS, Oxide). The cell pellets were suspended in 2.5% glutaraldehyde and stored at 4°C for 4 hr. Samples were rinsed three times with PBS, post-fixed in 1% osmium tetroxide for 1 hr, washed three times with PBS, dehydrated in ethanol solutions (60, 70, 80, 90% and twice at 100%) for 5 min, with 5 min intervals between treatments. The dehydrated samples were treated with 1 ml hexamethyldisilizine, dried overnight and coated with gold-palladium in a SEM coating unit E5000 (Polaron Equipment Ltd.) for observation under scanning electron microscope (JEOL JSM-6380LV)^42^.

### Transition of bacterial shape under nutrient stress

To determine the primary effect of nutrient concentration on cell morphology and growth, the strain YC6860^T^ was first cultured in 0.5 TSB for 15 days as described above and the same cell pellets were harvested by centrifugation (8000*g* at 4°C) for 10 min and washed twice with 0.1 TSB. The washed cells were transferred to 0.1 TSB and cultivated until 22 days after the initial inoculation of the strain. Samples of bacterial cells were collected at 7 and 16 days after cell transfer for observation of cell morphology and growth, and the measurement of total organic carbon (TOC) concentration. Harvest and transfer were performed aseptically to prevent contamination of bacterial cells during the transfer.

### Measurement of surface area-to-volume ratio of bacterial cells

To measure the S/V ratio of bacterial cells cultured in TSB containing various nutrient concentrations (0.1~0.5 TSB) for 15 days at 28°C, serial sectioning and reconstruction of 3D models using TEM were carried out as described previously^43^. In brief, sample blocks for TEM were sliced into serial section ribbons (120 nm thickness) and stained with uranyl acetate solution (3 min) and lead citrate solution (6 min). Images from serial sections were stacked into MRC format files and bacterial cells were outlined to prepare 3D models using the IMOD software package. Morphometric information of bacterial cells was acquired with the imodinfo command in the package and the S/V ratios were calculated using the Microsoft Excel program. In each sample, at least 8 bacterial cells were randomly selected to calculate their volume and surface. This experiment was repeated to confirm the changes in the S/V ratio. The relationship between TSB concentrations and S/V ratios of bacterial cells was subjected to regression analysis.

### Determination of total organic carbon

The TOC concentration in the culture supernatant of strain YC6860^T^ was determined via the 680°C combustion-infrared method using a Shimadzu carbon analyzer (TOC-VCPN, Japan). The carbon dioxide generated by oxidation was detected using infrared gas analyzer according to the manufacturer’s manual. The culture supernatant was prepared by centrifugation (5000*g*, 15 min) of bacterial cells cultured for 22 days at 28°C. Triplicates of each sample were analyzed.

### Phylogenetic analysis of 16S rRNA gene sequence

The genomic DNA was extracted from the strain YC6860^T^ using a commercial DNA extraction kit (Core Biosystem, Korea). The 16S rRNA gene was PCR amplified from purified genomic DNA using a set of primers 27F and 1492R^44^ and the purified PCR product was sequenced by Geno Tech Inc. (Daejeon, Korea). The 16S rRNA gene sequences were compiled using SeqMan software (DNASTAR) and the sequences of related taxa were obtained from the GenBank database. The chimera in the sequence of strain was checked using the CHECK_CHIMERA program in the Ribosomal Database Project (RDP)^45^. The phylogenetic position of strain YC6860^T^ was determined by comparing its 16S rRNA gene sequence and sequences of related taxa collected from NCBI and EzBioCloud server (http://www.ezbiocloud.net)^46^. Multiple alignments were performed using CLUSTAL_X program^47^ and a BioEdit program was used for gaps editing^48^. The evolutionary distances were calculated using the Kimura two-parameter model^49^. The phylogenetic tree was constructed using a neighbor-joining method^50^ and maximum-parsimony^51^ in a MEGA4 program^52^ with boot strap values based on 1000 replications^53^.

### Physiological and biochemical tests

API ZYM, API 20E and API 20NE kits were used for physiological and biochemical characterization of strain YC6860^T^. API ZYM strips were read after 5-hr incubation. The assimilation of single substrate was determined using API 20NE kits at 30°C after 24-hr incubation. The bacterial cells were grown at different temperatures ranging from 4°C-40°C. Anaerobic growth was tested at 28°C by pouring a thick layer of vaspar (50% petrolatum, 50% paraffin) on the surface of inoculated semi-solid half-strength R2A agar (1.0% agar) in 35 mL screw capped glass tubes^54^. Growth in sodium chloride was observed at 0-5% (w/v) while growth at pH 5-10 was determined after 7 days of incubation. Oxidase and catalase activities were determined according to the method of Cappuccino & Sherman^55^. Gram staining was carried out using a gram staining kit (BD) according to the manufacturer’s instructions.

### Chemotaxonomic characterization

The G+C content of the chromosomal DNA was determined after purification and extraction of genomic DNA as previously reported^56^. It was then degraded enzymatically into nucleosides and the G+C contents were determined via reverse phase HPLC^57^. The quinone system was determined by TLC (Tindall, 1990). The fatty acid profile was determined by incubating the strain YC6860^T^ either in R2A or 0.1 TSB and 0.5 TSB at 28°C for 10 days. According to the standard protocol of Sasser (1990), the cellular fatty acids were saponified, methylated and extracted and the composition was determined using GC (6890; Hewlett Packard) and a microbial identification software package (Microbial ID).

### Full genome sequence of strain YC6860^T^

Complete genome of strain YC6860^T^ was sequenced by extracting its high-quality genomic DNA with the blood and cell culture DNA minikit (Qiagen, MD). The genome of YC6860^T^ was sequenced and characterized using Illumina GAIIx sequencer (San Diego, CA) with 100-bp paired-end libraries from Chun-Lab, Inc. (Seoul, Korea). GS Assembler 2.3 (Roche Diagnostics, Branford, CT) and Codon Code Aligner 3.0 were used to assemble all sequence data and the gaps between all the contigs were filled by PCR amplification. The coding sequences (CDSs) were determined with Glimmer 3.02. Transfer RNAs (tRNAs) were determined by tRNAscan-SE^58^, while the ribosomal RNA (rRNA) was located using HMMER with EzBiocloud-e-rRNA profiles^46,59^. The CDSs were compared by rpsBLAST and NCBI reference sequences (Ref-Seq) to catalytic families (Catfam) and the NCBI COG. The functional annotation of the CDSs was determined by employing the SEED database^60,61^.

### Accession number of 16S rRNA gene and genome sequence of YC6860^T^

The nucleotide sequence of 16S rRNA gene and the full genome sequence of the strain YC6860^T^ were deposited under the accession numbers of GQ369128 and CP007440, respectively.

## Acknowledgements

This work was carried out with the support of “Cooperative Research Program for Agriculture Science & Technology Development (PJ 01104901, PJ 010953092018)” Rural Development Administration, Republic of Korea. E.J. Chung and A. Khan were supported by a scholarship awarded by the BK21 Plus Program, the Ministry of Education, Republic of Korea.

## SUPPLEMENTAL MATERIALS

**Table S1.**
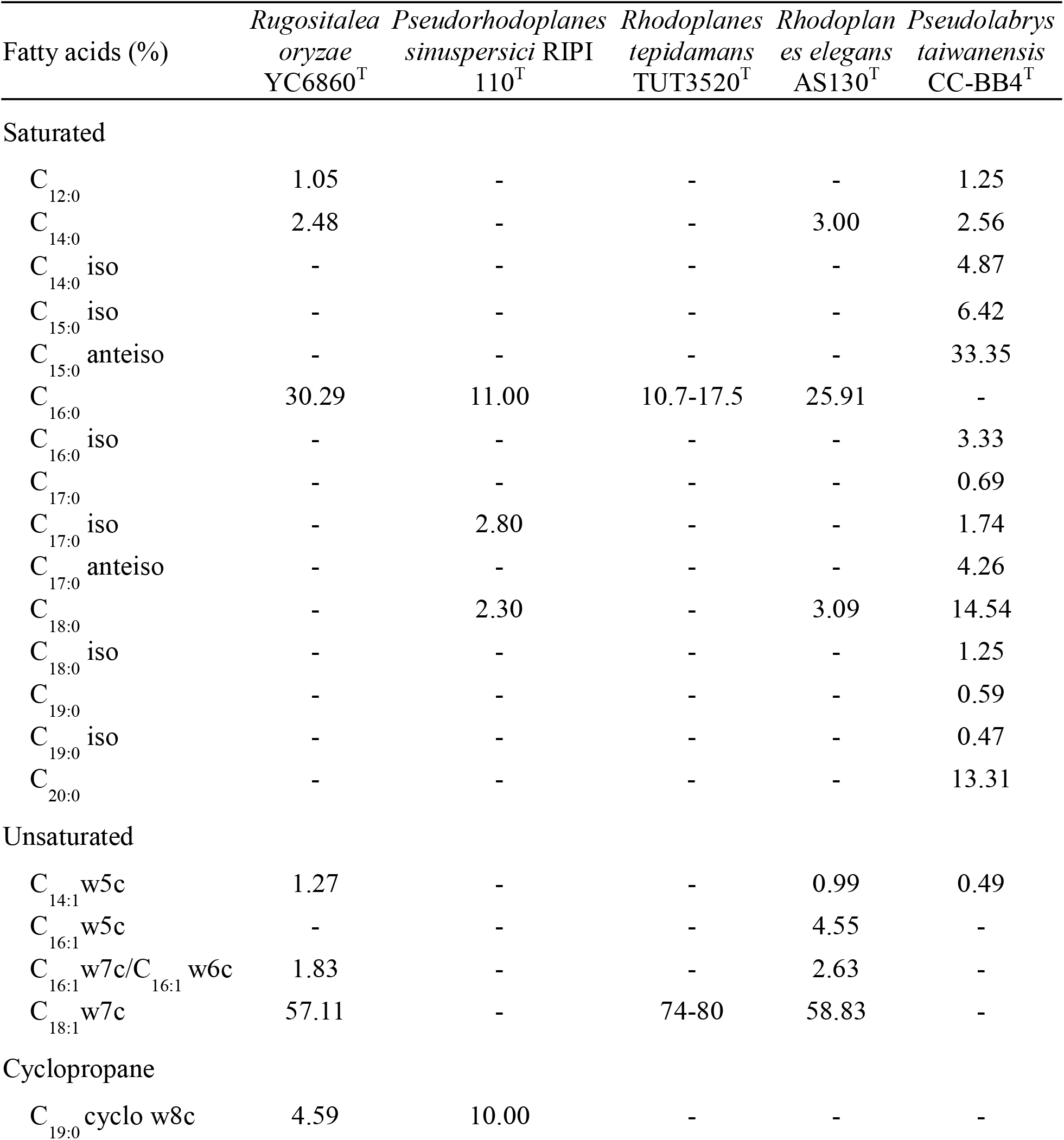
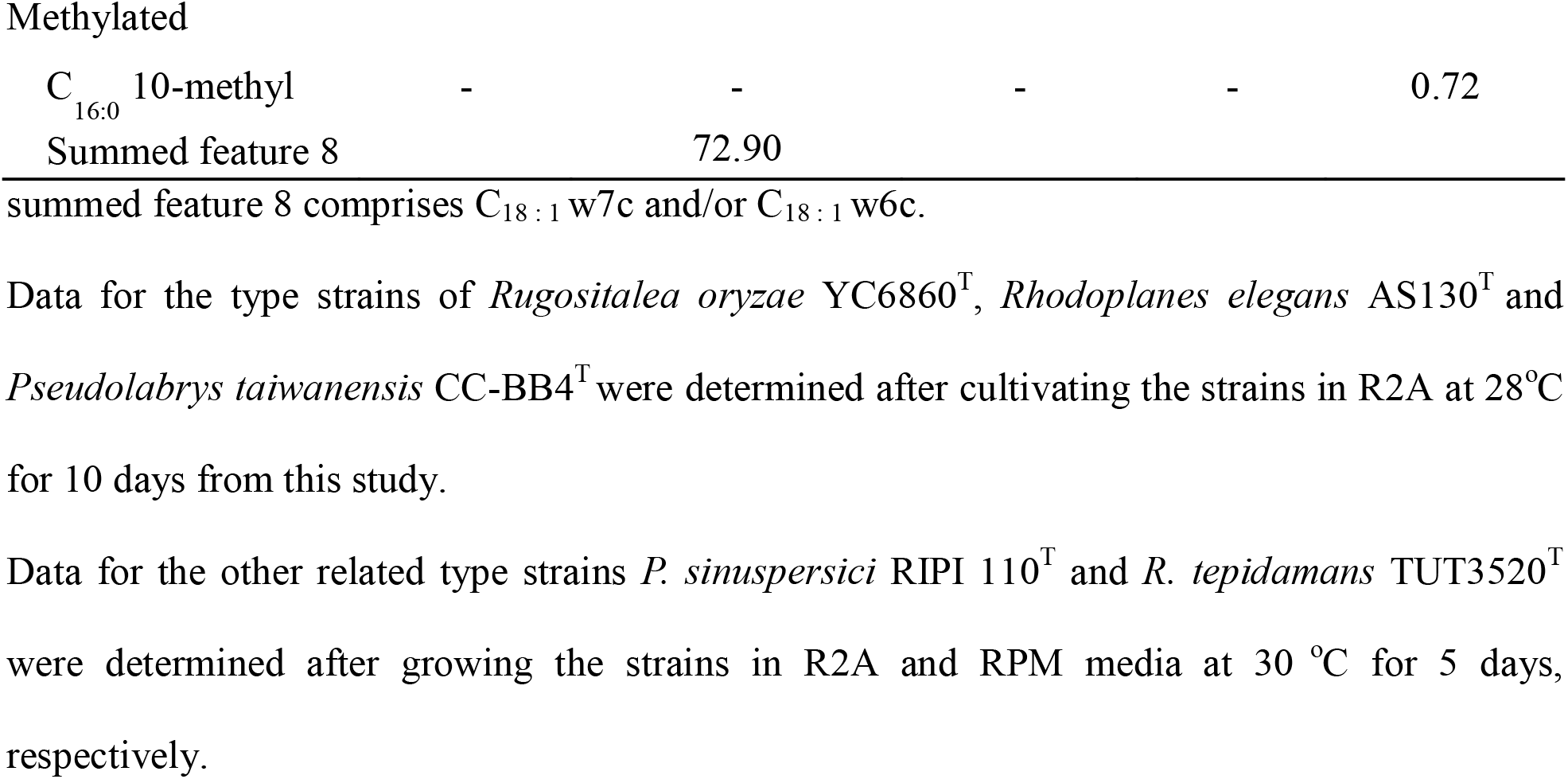
Cellular fatty acid profiles of *Rugositalea oryzae* YC6860^T^ and phylogenetically related genera in the order *Rhizobiales*. Taxa: *Rugositalea oryzae* YC6860^T^; *Pseudorhodoplanes sinuspersici* RIPI 110^T^; *Rhodoplanes tepidamans* TUT3520^T^; *Rhodoplanes elegans* AS130^T^; *Pseudolabrys taiwanensis* CC-BB4^T^.

**Table S2.**
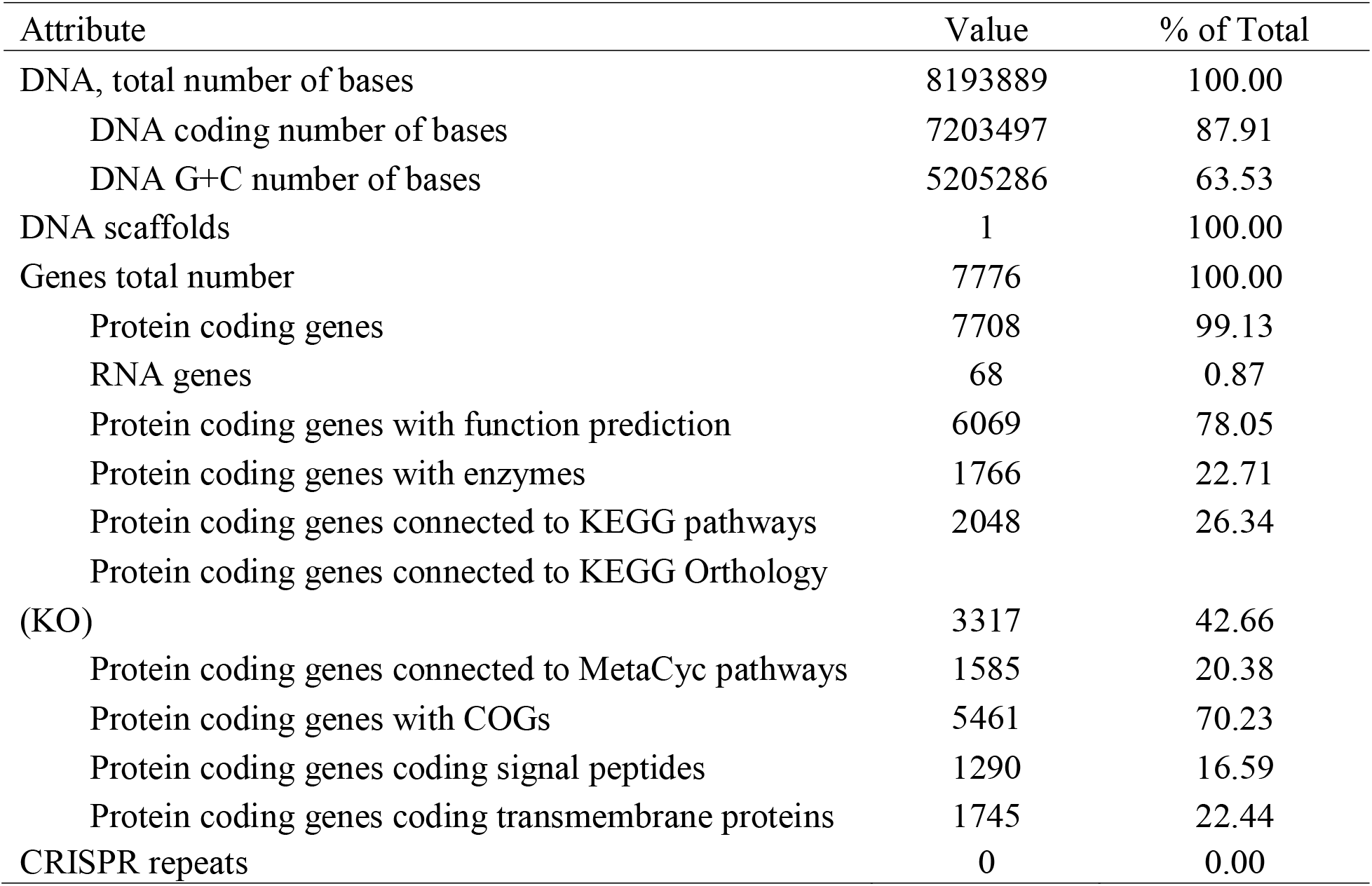
Nucleotide content and gene count levels of the *Rugositalea oryzae* YC6860^T^ genome.

**Figure S1.**
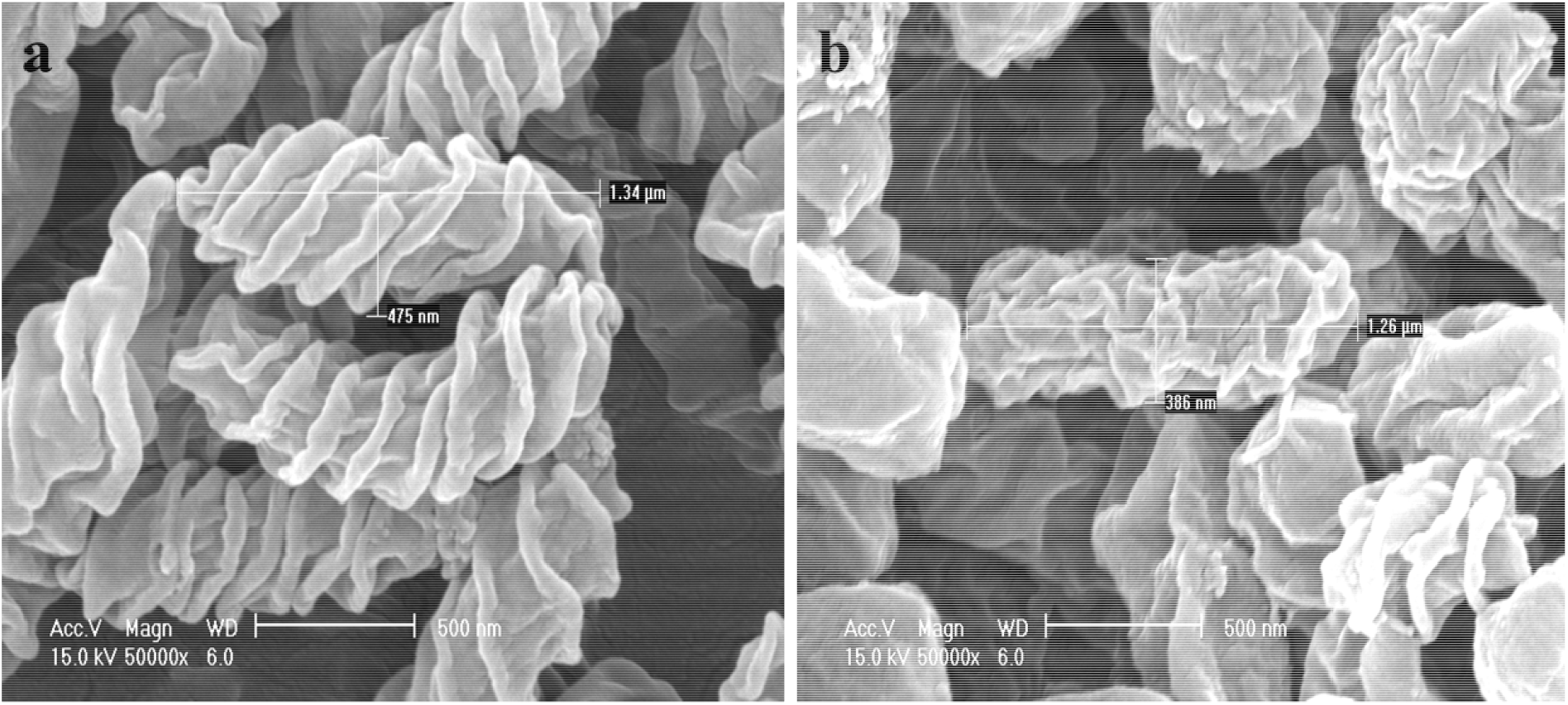
The growth of *Rugositalea oryzae* YC6860^T^ cells grown on (a) 0.1 LB and (b) 0.5 LB broth at 28°C in a rotary shaker (50 rpm) for 5 days.

**Figure S2.**
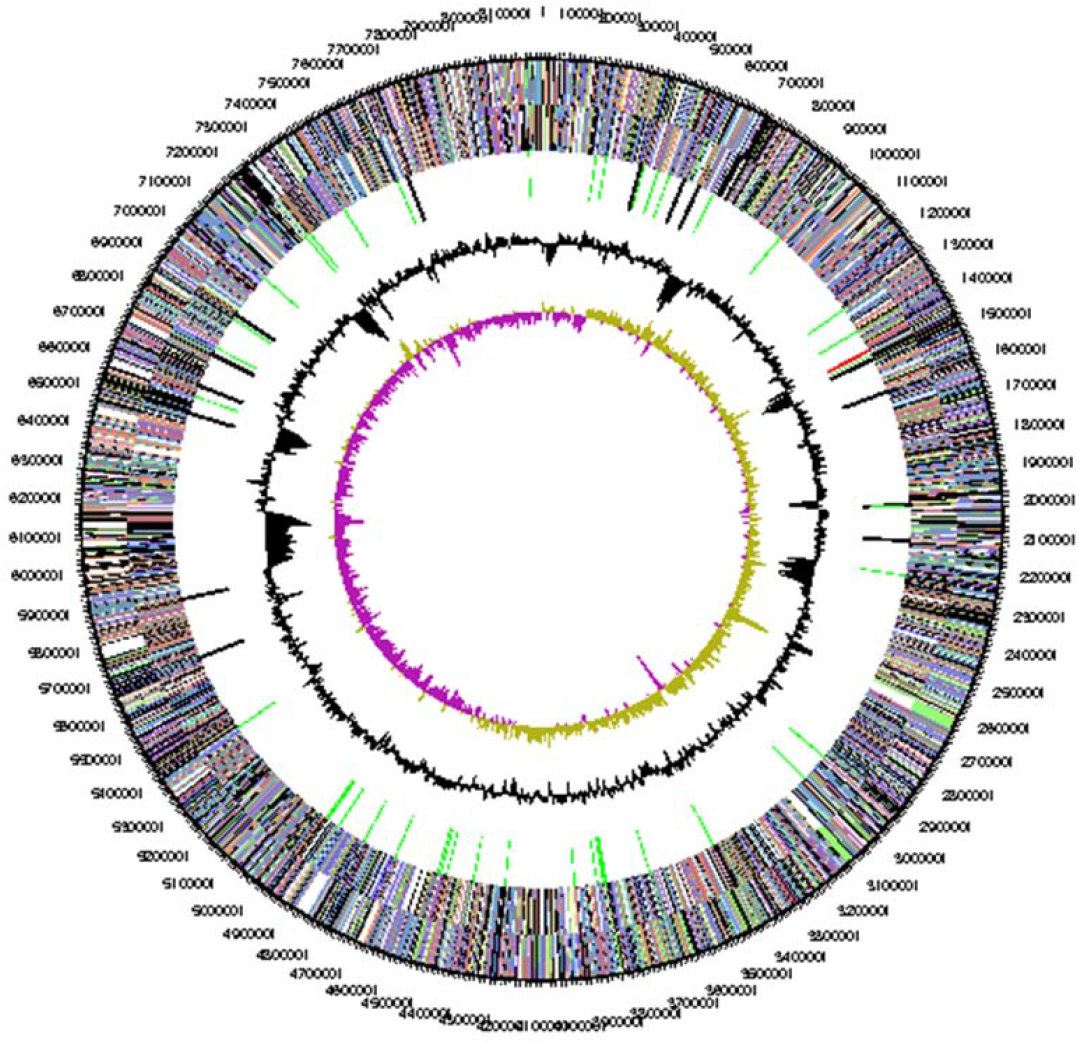
Circular representation of the *Rugositalea oryzae* YC6860^T^ genome. Circles from th outside to the center: genes on forward strand colored by COGs categories, genes on reverse strand (colored by COG categories), tRNA (green), rRNA (red), other RNAs (black), GC content and GC skew.

**Figure S3.**
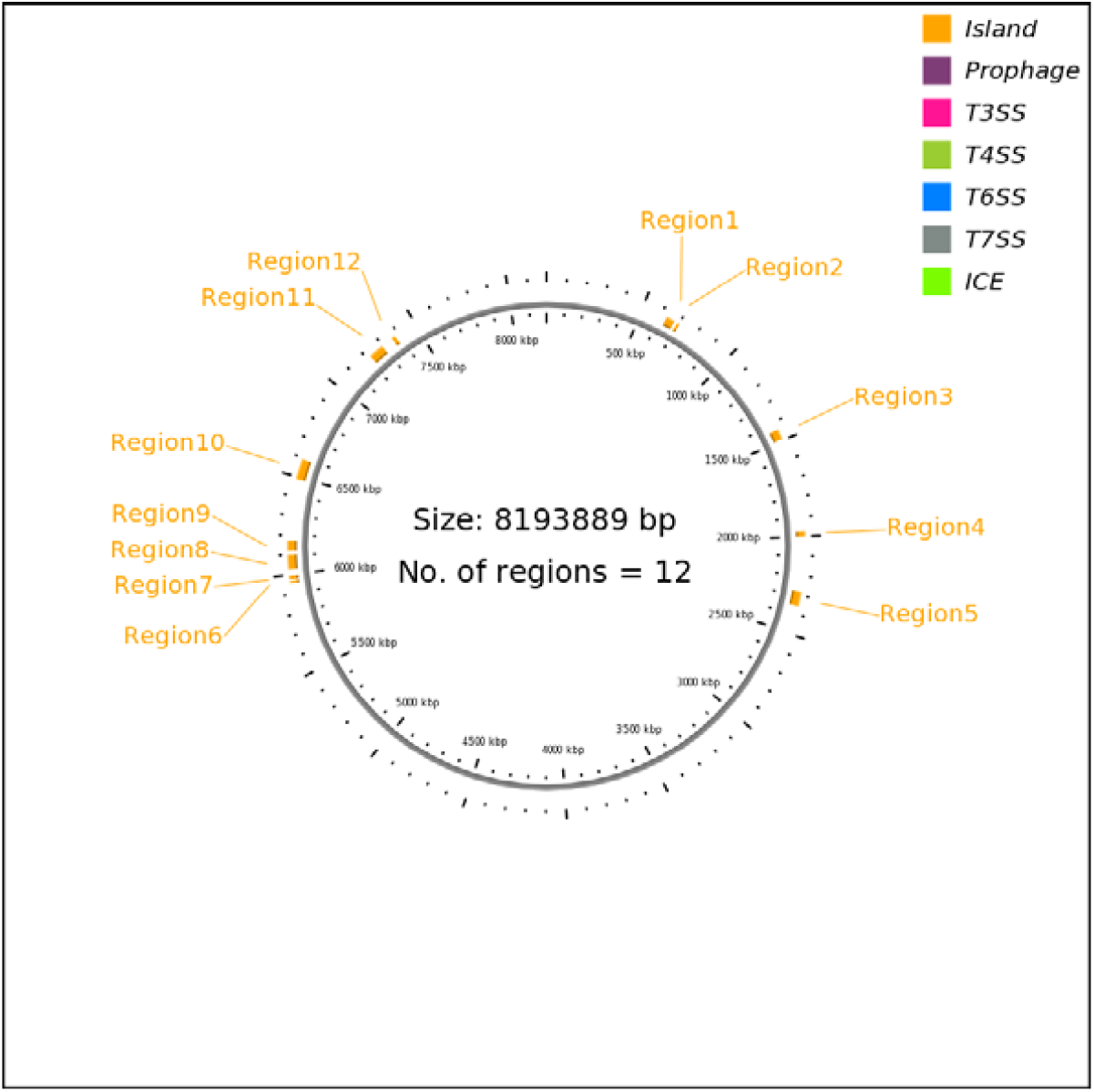
Map of the island like regions in the *Rugositalea oryzae* YC6860^T^ genome.

**Figure S4.**
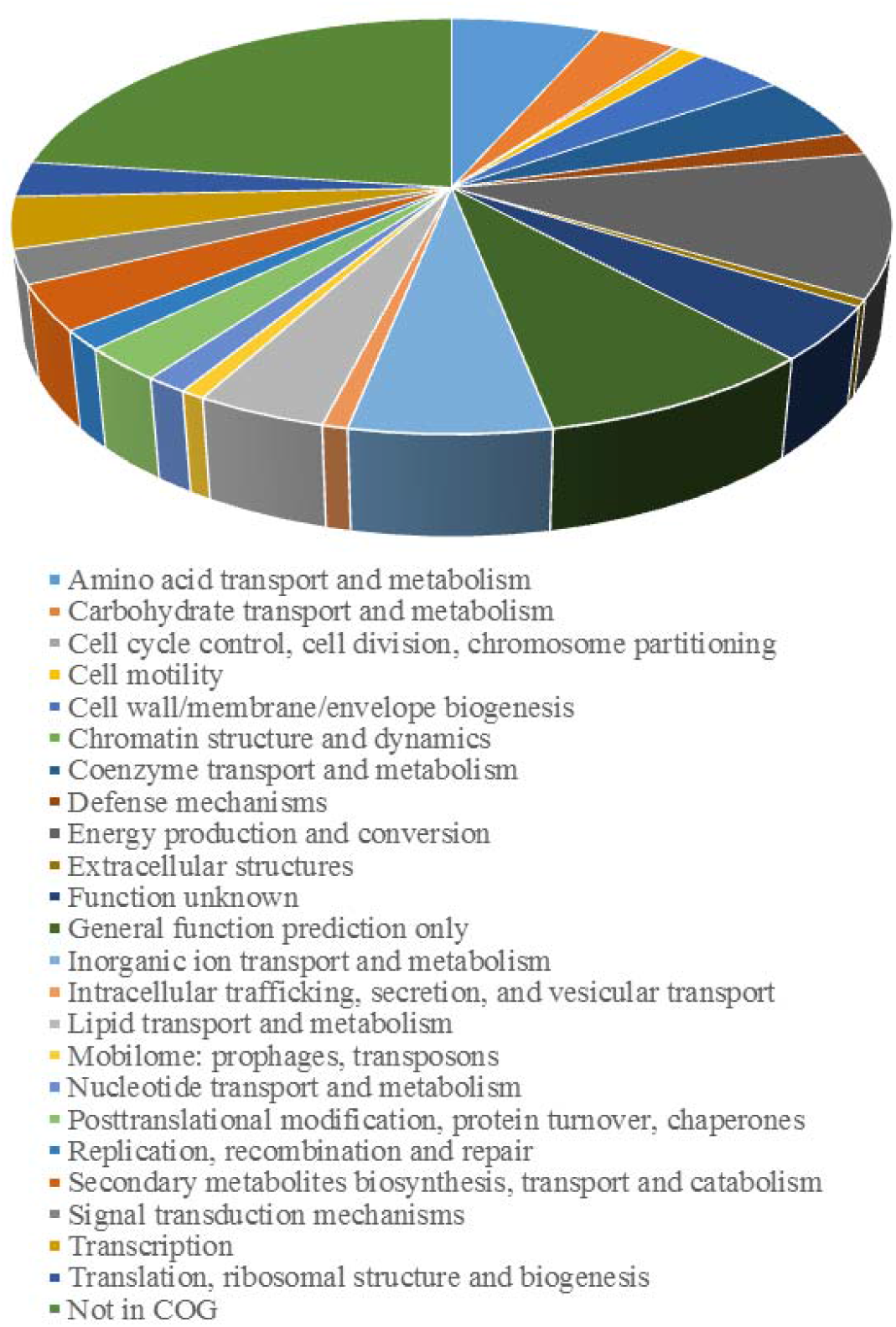
The distribution of genes into COGs functional categories of the *Rugositalea oryzae* YC6860^T^ genome.

**Figure S5.**
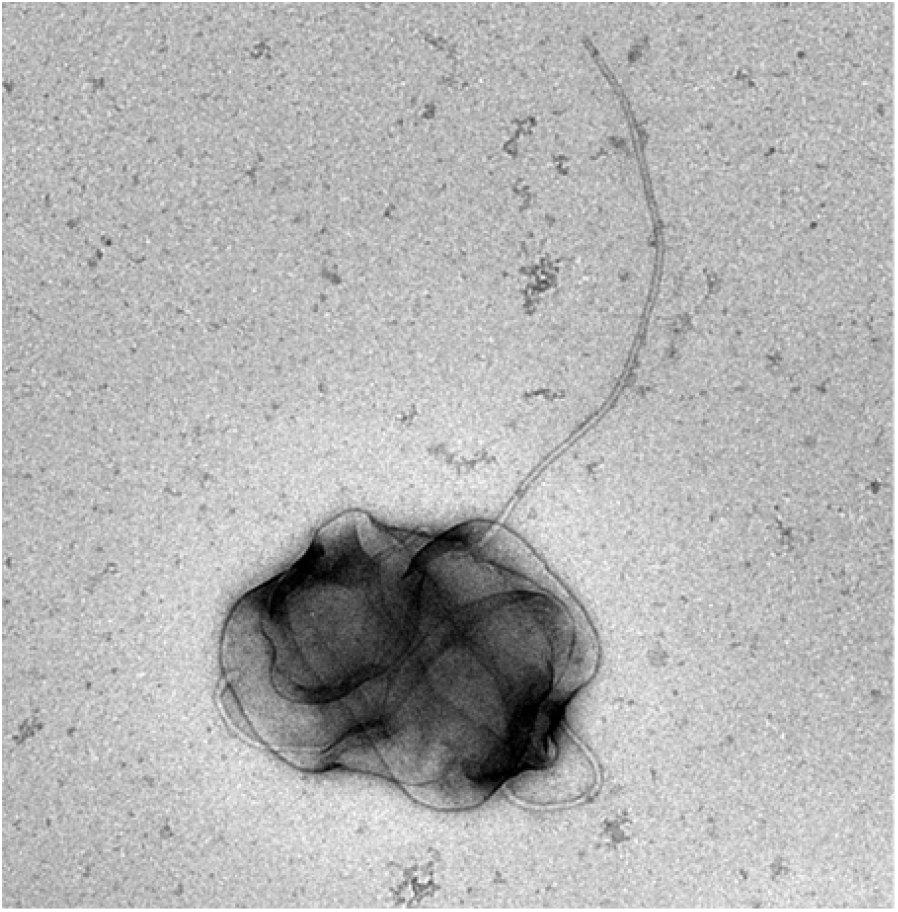
Cells of *Rugositalea oryzae* YC6860^T^ with a polar flagellum grown in 0.1 TSB at 28°C in a rotary shaker (50 rpm) for 5 days.

## Notes

### Competing Interest Statement

The authors have declared no competing interest.

