## Supplementary information for "Unusual Morphological Changes of *Rugositalea oryzae*, A Novel Wrinkled Bacterium Isolated from The Rice Rhizosphere, Under Nutrient Stress"

**Table S1.** Cellular fatty acid profiles of *Rugositalea oryzae* YC6860^T^ and phylogenetically related genera in the order *Rhizobiales.* Taxa: *Rugositalea oryzae* YC6860^T^; *Pseudorhodoplanes sinuspersici* RIPI 110^T^; *Rhodoplanes tepidamans* TUT3520^T^; *Rhodoplanes elegans* AS130^T^; *Pseudolabrys taiwanensis* CC-BB4^T^.

| Fatty acids (%) | *Rugositalea oryzae* YC6860^T^ | *Pseudorhodoplanes sinuspersici* RIPI 110^T^ | *Rhodoplanes tepidamans* TUT3520^T^ | *Rhodoplanes elegans* AS130^T^ | *Pseudolabrys taiwanensis* CC-BB4^T^ |
| --- | --- | --- | --- | --- | --- |
| Saturated |  |  |  |  |  |
| C_12:0_ | 1.05 | - | - | - | 1.25 |
| C_14:0_ | 2.48 | - | - | 3.00 | 2.56 |
| C_14:0_ iso | - | - | - | - | 4.87 |
| C_15:0_ iso | - | - | - | - | 6.42 |
| C_15:0_ anteiso | - | - | - | - | 33.35 |
| C_16:0_ | 30.29 | 11.00 | 10.7-17.5 | 25.91 | - |
| C_16:0_ iso | - | - | - | - | 3.33 |
| C_17:0_ | - | - | - | - | 0.69 |
| C_17:0_ iso | - | 2.80 | - | - | 1.74 |
| C_17:0_ anteiso | - | - | - | - | 4.26 |
| C_18:0_ | - | 2.30 | - | 3.09 | 14.54 |
| C_18:0_ iso | - | - | - | - | 1.25 |
| C_19:0_ | - | - | - | - | 0.59 |
| C_19:0_ iso | - | - | - | - | 0.47 |
| C_20:0_ | - | - | - | - | 13.31 |
| Unsaturated |  |  |  |  |  |
| C_14:1_w5c | 1.27 | - | - | 0.99 | 0.49 |
| C_16:1_w5c | - | - | - | 4.55 | - |
| C_16:1_w7c/C_16:1_ w6c | 1.83 | - | - | 2.63 | - |
| C_18:1_w7c | 57.11 | - | 74-80 | 58.83 | - |
| Cyclopropane |  |  |  |  |  |
| C_19:0_ cyclo w8c | 4.59 | 10.00 | - | - | - |
| Methylated |  |  |  |  |  |
| C_16:0_ 10-methyl | - | - | - | - | 0.72 |
| Summed feature 8 |  | 72.90 |  |  |  |

summed feature 8 comprises C_18 : 1_ w7c and/or C_18 : 1_ w6c.

Data for the type strains of *Rugositalea oryzae* YC6860^T^, *Rhodoplanes elegans* AS130^T^ and *Pseudolabrys taiwanensis* CC-BB4^T^ were determined after cultivating the strains in R2A at 28^o^C for 10 days from this study.

Data for the other related type strains *P. sinuspersici* RIPI 110^T^ and *R. tepidamans* TUT3520^T^ were determined after growing the strains in R2A and RPM media at 30 ^o^C for 5 days, respectively.

**Table S2.** Nucleotide content and gene count levels of the *Rugositalea oryzae* YC6860^T^ genome.

| Attribute | Value | % of Total |
| --- | --- | --- |
| DNA, total number of bases | 8193889 | 100.00 |
| DNA coding number of bases | 7203497 | 87.91 |
| DNA G+C number of bases | 5205286 | 63.53 |
| DNA scaffolds | 1 | 100.00 |
| Genes total number | 7776 | 100.00 |
| Protein coding genes | 7708 | 99.13 |
| RNA genes | 68 | 0.87 |
| Protein coding genes with function prediction | 6069 | 78.05 |
| Protein coding genes with enzymes | 1766 | 22.71 |
| Protein coding genes connected to KEGG pathways | 2048 | 26.34 |
| Protein coding genes connected to KEGG Orthology (KO) | 3317 | 42.66 |
| Protein coding genes connected to MetaCyc pathways | 1585 | 20.38 |
| Protein coding genes with COGs | 5461 | 70.23 |
| Protein coding genes coding signal peptides | 1290 | 16.59 |
| Protein coding genes coding transmembrane proteins | 1745 | 22.44 |
| CRISPR repeats | 0 | 0.00 |

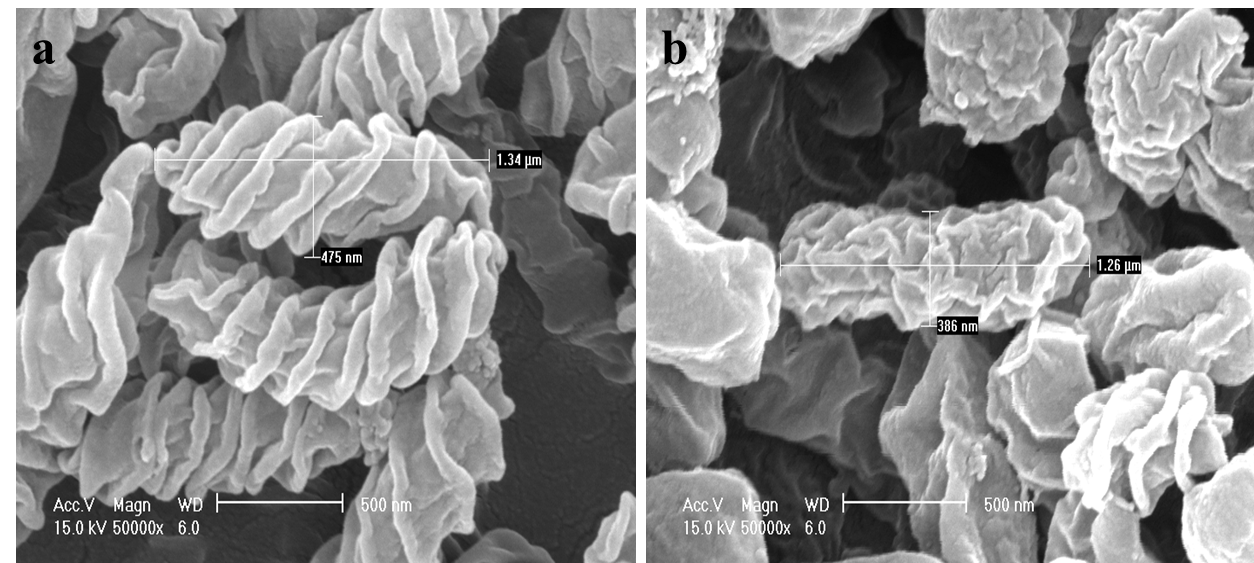

**Figure S1.** The growth of *Rugositalea oryzae* YC6860^T^ cells grown on (a) 0.1 LB and (b) 0.5 LB broth at 28^o^C in a rotary shaker (50 rpm) for 5 days.

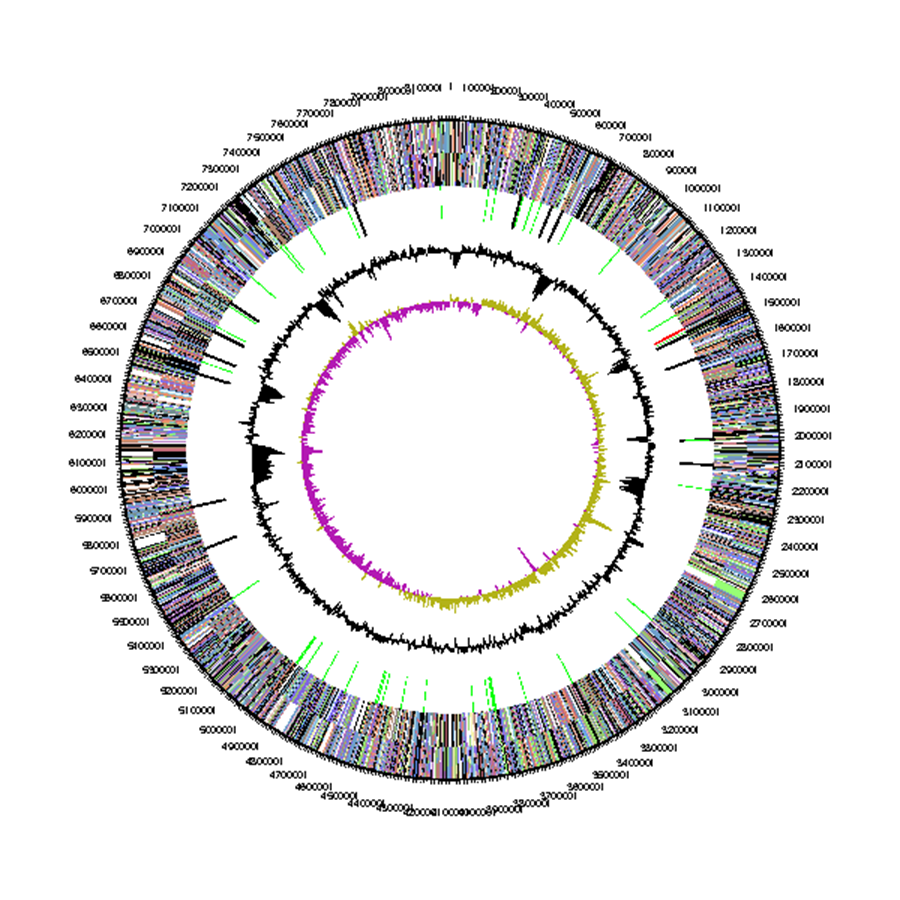

**Figure S2.** Circular representation of the *Rugositalea oryzae* YC6860^T^ genome. Circles from the outside to the center: genes on forward strand colored by COGs categories, genes on reverse strand (colored by COG categories), tRNA (green), rRNA (red), other RNAs (black), GC content and GC skew.

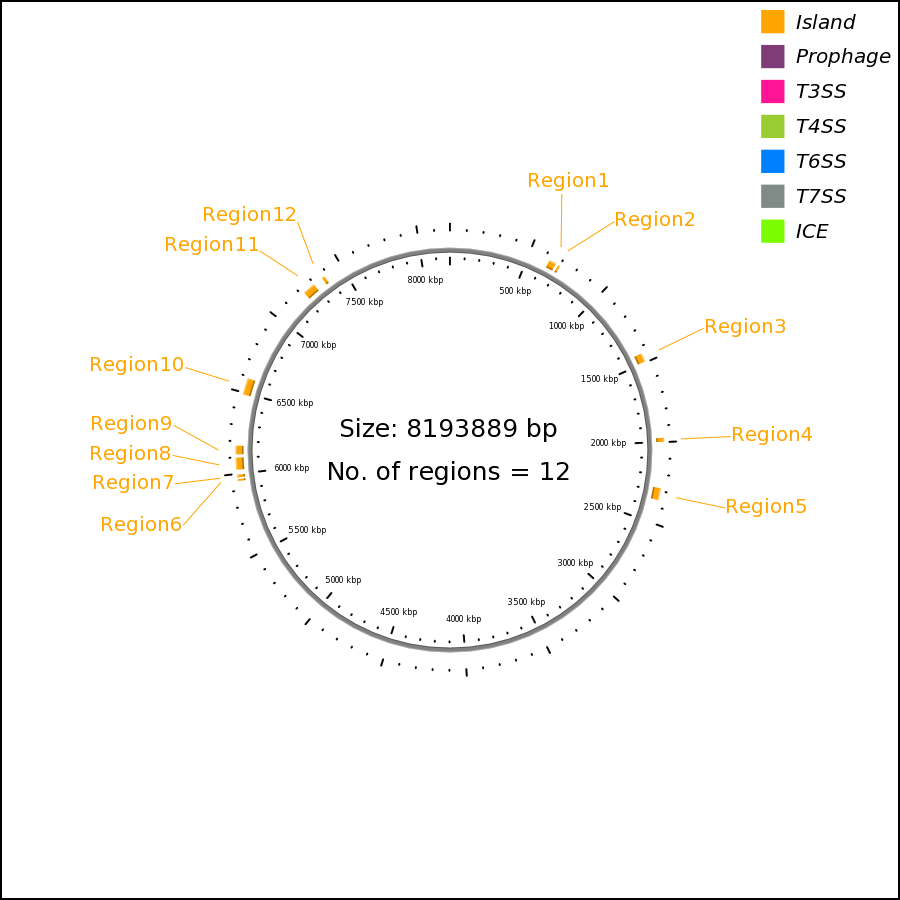

**Figure S3.** Map of the island like regions in the *Rugositalea oryzae* YC6860^T^ genome.

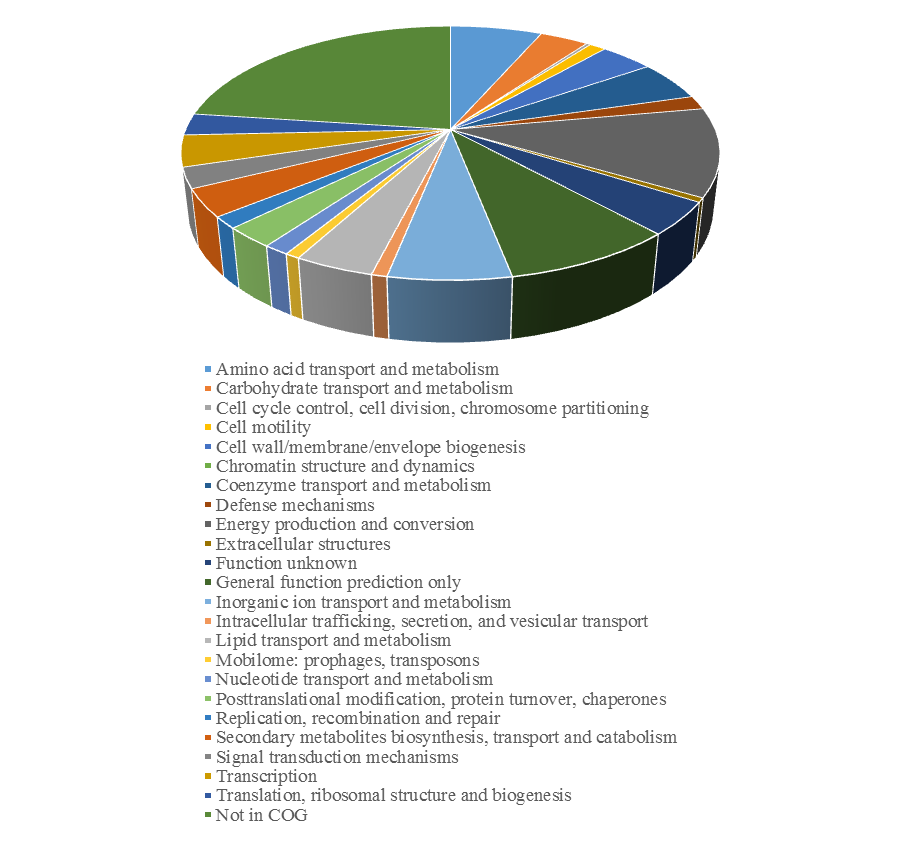

**Figure S4.** The distribution of genes into COGs functional categories of the *Rugositalea oryzae* YC6860^T^ genome.

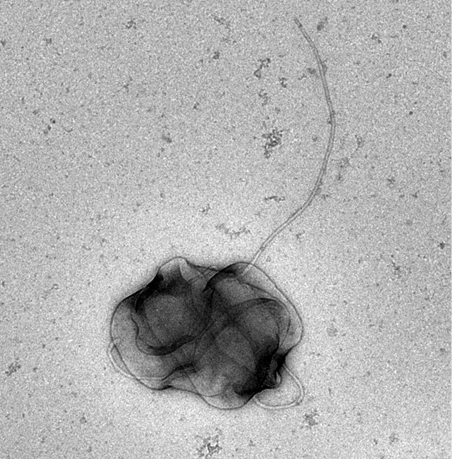

**Figure S5.** Cells of *Rugositalea oryzae* YC6860^T^ with a polar flagellum grown in 0.1 TSB at 28^o^C in a rotary shaker (50 rpm) for 5 days.
